# Parameter estimation in a whole-brain network model of epilepsy: comparison of parallel global optimization solvers

**DOI:** 10.1101/2023.11.01.565083

**Authors:** David R. Penas, Meysam Hashemi, Viktor K. Jirsa, Julio R. Banga

## Abstract

The Virtual Epileptic Patient (VEP) refers to a computer-based representation of a patient with epilepsy that combines personalized anatomical data with dynamical models of abnormal brain activities. It is capable of generating spatio-temporal seizure patterns that resemble those recorded with invasive methods such as stereoelectro EEG data, allowing for the evaluation of clinical hypotheses before planning surgery. This study highlights the effectiveness of calibrating VEP models using a global optimization approach. The approach utilizes SaCeSS, a cooperative metaheuristic capable of parallel computation, to yield high-quality solutions without requiring excessive computational time. Through extensive benchmarking, our proposal successfully solved a set of different configurations of VEP models, demonstrating better scalability and superior performance against other parallel solvers. These results were further enhanced using a Bayesian optimization framework for hyperparameter tuning, with significant gains in terms of both accuracy and computational cost. Additionally, we added an scalable uncertainty quantification phase after model calibration, and used it to assess the variability in estimated parameters across different problems. Overall, this study has the potential to improve the estimation of pathological brain areas in drug-resistant epilepsy, thereby to inform the clinical decision-making process.

**Author summary:** Motivated by the problem of parameter estimation in a set of whole-brain network models of epilepsy (of increasing complexity), this study addresses the question of choosing a robust global optimization solver that can be accelerated by exploiting parallelism in different infrastructures, from desktop workstations to supercomputers. By leveraging data-driven techniques with robust cooperative global optimization methods, we aim to achieve accurate parameter estimation with reduced reliance on prior information. This is due to the dependency of Bayesian inference on the level of information in the prior, while this approach allows us to quantify uncertainty in the absence of any prior knowledge effectively. In this work, we construct an efficient and accurate method to perform parameter estimation and uncertainty quantification for the VEP model, and we use it to infer the brain regional epileptogenicity from source and sensor level whole-brain data. Of specific interest is the ability of our method to produce inference for high-dimensional state-space models governed by deterministic, stochastic, well-behaved, and stiff differential equations, using only partial observations and sparse encoding from system states to the observation.

## Introduction

Adjusting the stimulation parameters poses a significant challenge in inferring critical information, such as characterizing pathological brain networks. The ability to precisely manipulate the stimulation parameters is essential for gaining insights into underlying mechanisms of the data-generating process in pathological conditions, such as the dynamics and connectivity of abnormal brain networks.

In partial epilepsy, seizures originate from a network of hyperexcitable regions referred to as epileptogenic zone (EZ; [1]) and then propagate to a secondary connected network, the so-called propagation zone (PZ; [2]). The success of surgical interventions for drug-resistant patients critically depends on the precision and reliability of the initial hypotheses, e.g., the spatial map of EZ/PZ as an identification of the seizure organization [3–8].

Virtual Epileptic Patient (VEP; [7, 9]) is a digital modeling approach that integrates mathematical modeling of abnormal neural activity with patient-specific anatomical data to predict the brain network involved in seizure generation and propagation in individuals. By accurately capturing diverse seizure dynamics and generating computer simulations resembling intracranial EEG recordings, this technique offers a versatile platform to optimize the surgical strategy and to aid in clinical decision-making [7, 8, 10, 11]. The VEP is a model-based approach that relies on estimating the parameters in a high-dimensional state-space representation to accurately identify the network of EZ/PZ in the brain. Besides the issues of sparsity, stochasticity, and scalability, the reliability of prediction on the EZ/PZ is challenging due to the nontrivial effects of brain networks, the non-linearity involved in the spatiotemporal organization of the brain, and the uncertainty in model components.

Several studies [5, 7, 8, 10, 12–15] have demonstrated that advancements in more accurate estimation of the VEP parameters can lead to more informed clinical decision-making and optimize surgical strategies. This motivation drove us to benchmark various parallel global search optimization techniques that eliminate the need for strong initial assumptions, such as informative prior information in a Bayesian setup [5]. Global search optimization methods offer several advantages, such as an scalability and flexibility in objective functions and constraints, enhanced exploration-exploitation trade-off, robustness to initial conditions, and the capability to deal with non-convex and multimodal problems. By leveraging the advantages of global search optimization methods with high-performance computing (HPC) infrastructures, we can achieve faster and more accurate parameter estimation, thereby enhancing the overall efficacy of our approach.

## Materials and methods

### Network anatomy

The structural connectome was built using the TVB-specific reconstruction pipeline with generally available neuroimaging software, as described in [10, 16]. First, the command *recon-all* from the Freesurfer package [17] in version v6.0.0 was used to reconstruct and parcellate the brain anatomy from T1-weighted images. Then, the T1-weighted images were coregistered with the diffusion-weighted images using the linear registration tool *flirt* [18] from the FSL package in version 6.0, with the correlation ratio cost function and 12 degrees of freedom.

The MRtrix package version 0.3.15 was then used for tractography. The fiber orientation distributions were estimated from diffusion-weighted images using spherical deconvolution [19] by the *dwi2fod* tool, with the response function estimated by the *dwi2response* tool using the *tournier* algorithm [20]. Next, we used the *tckgen* tool, employing the probabilistic tractography algorithm iFOD2 [21] to generate 15 million fiber tracts. Finally, with the generated fiber tracts and the regions defined by the brain parcellation, the connectome matrix is built by counting the fibers connecting all regions. Using the *tck2connectome* tool and the Desikan-Killiany parcellation [22], the patient’s brain is divided into 68 cortical regions and 16 subcortical structures. See Table S2 for the label names and indices of the sub-divided brain regions. The connectome was normalized so that the maximum value is equal to one (see S1 Fig).

### Virtual Epileptic Patient (VEP) model

In the process of building a personalized brain network model, the brain regions (network nodes) are defined using a parcellation scheme, and a set of dynamical equations, known as the neural mass model, are placed at each network node to generate the regional brain activity [9, 23]. Neural masses are commonly used to model the collective behavior of populations of neurons in the brain, such as firing rates, capturing macroscopic dynamics and interactions rather than focusing on individual neuron behavior [24–27]. They have demonstrated efficiency in capturing the main features of brain functional behaviors in a single computational framework by accounting for interactions among brain regions in healthy and diverse pathological conditions [7, 14, 23, 28–31]. Taking a data-driven approach that integrates subject-specific brain anatomy, the network edges are subsequently encoded using a personalized structural connectivity matrix derived from non-invasive imaging data, such as diffusion magnetic resonance imaging (dMRI), for an individual subject [9, 10]. Moreover, the anatomical data imposes a constraint on simulated brain network dynamics, that is, the evolution of trajectories in the latent space, allowing the hidden state dynamics to be inferred from the data. In the Virtual Epileptic Patient (VEP model; [9]), a personalized brain network model of epilepsy spread, the dynamics of brain network nodes are governed by the so-called Epileptor model [32]. The Epileptor is a general description of epileptic seizures (in humans, mice, rats, and zebrafishes), which contains the complete taxonomy of system bifurcations to realistically reproduce the dynamics of onset, progression, and offset of seizure-like events [33, 34]. The full Epileptor model consists of five state variables that couple two oscillatory dynamical systems operating on three distinct time scales. At the fastest time scale, an oscillatory dynamical system accounts for fast discharges during ictal seizure states, whereas on the intermediate time scale, another system represents the slow spike-and-wave oscillations. On the slowest time scale, a permittivity state variable is responsible for the transition between interictal and ictal states. The permittivity variable represents the slow-evolving extracellular processes that occur during epileptiform activity, such as levels of ions, oxygen, and energy metabolism. Depending on its values, the dynamics of Epileptor may drive it into or out of a seizure, which accounts for its bi-stable behavior (for more details, see [32]).

Motivated by Synergetic theory [35, 36] and assuming a time-scale separation (τ ≫ 1), the fast variables swiftly converge onto the slow manifold, which is governed by the dynamics of the slow variable. This adiabatic approximation [37, 38] leads to the 2D reduction of VEP model, given by:

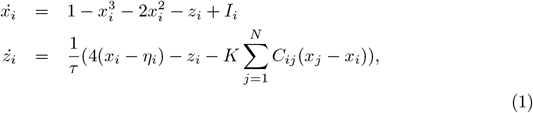

where *x*_*i*_ and *z*_*i*_ indicate the fast and slow variables corresponding to *i*-th brain region, respectively. The parameter *I* = 3.1 represents the flow of electric current, and τ scales the length of the seizure. The degree of epileptogenicity at each brain region is represented by the value of the excitability parameter η_*i*_ (Hopf bifurcation parameter). The network nodes are coupled by a linear diffuse approximation of permittivity coupling through 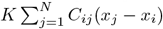, which includes a global scaling factor *K* on the subject’s connectome *C*_*ij*_.

### Spatial Map of Epileptogenicity

In the VEP model, the occurrence of seizures in each brain region depends on its individual excitability (node dynamics) and its connectivity to other regions (network coupling). The excitability parameter, denoted by η, regulates the level of tissue excitability, and its spatial distribution is the focus of parameter estimation. The brain regions can be classified into three main types based on their excitability parameter:

- Epileptogenic Zone (EZ): If η > η_*c*_, the brain region will autonomously trigger seizures, which are responsible for initiating and organizing the early stages of epileptic activity. The Epileptor model exhibits an unstable fixed point in these regions, allowing seizures to occur independently of network effects.
- Propagation Zone (PZ): If η_*c*_ − Δη < η < η_*c*_, the brain region does not autonomously trigger seizures. However, it can be recruited in seizure propagation since its equilibrium state is close to the critical value. In these regions, a supercritical Andronov–Hopf bifurcation occurs at η = η_*c*_, triggering seizure onset when a sufficiently large external input is present. Otherwise, the Epileptor model remains in a stable equilibrium state.
- Healthy Zone (HZ): If η < η_*c*_ − Δη, the brain region remains seizure-free. In these regions, all trajectories in the phase-plane of the Epileptor model converge to a single stable fixed point, indicating a healthy (non-epileptic) state.

Based on the above dynamical properties, the spatial distribution of epileptogenicity across different brain regions is determined by estimated heterogeneity in excitability parameters: EZs exhibiting high excitability, while PZs have lower excitability values, and very low values of excitability characterize all other regions as HZs.

Note that having an intermediate excitability value (i.e., close to the bifurcation value) does not guarantee recruitment into the seizure propagation. Seizure recruitment is governed by various factors, including structural connectivity, network coupling, and brain state dependence on noise, which all play a crucial role in determining the extent of propagation within the brain network [5, 10]. The linear stability analysis indicates that, in the absence of coupling (*K* = 0), the isolated nodes exhibit a bifurcation at the critical value η_*c*_ = −2.05 [39], and we set Δη = 1.0 [5, 10].

### Simulated Stereotactic-EEG (SEEG) data

Simulated Stereotactic-EEG (SEEG) implantation produces data to be used in the building and validation of VEP models. This invasive method is used in clinical situations for patients with drug-resistant epilepsy to determine the focal location of epileptic seizures [1, 7, 8, 40]. The implanted SEEG electrodes record the local field potential generated by the neuronal tissue in its vicinity. The gain matrix (also known as the lead-field or projection matrix) maps the source activity to the measurable sensor signals. Each sensor collects the source signals in its proximity, weighted by the distance and orientation of sources. To model the SEEG signals, here we assume an exponential relation between the source activities and the measurable signals at the sensors:

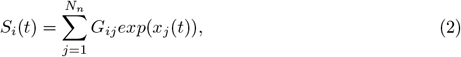

where *S*_*i*_(*t*) is the SEEG signal at sensor *i* ∈ {1, 2, …, *N*_*s*_} with *N*_*s*_ the number of channels (sensors), *x*_*j*_(t) is the source activity (given by fast variable in Eq. 1) in region *j* {1, 2, …, *N*_*n*_} with *N*_*n*_ the number of brain regions, and *G*_*ij*_ is the element of the gain matrix representing the distances of the sensors from the sources. Assuming that the generated signal decays with the square of the distance from the source, the gain matrix is approximated by

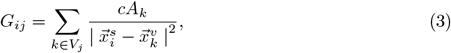

where *V*_*j*_ is the set of all vertices on the triangulated surface of region *j, c* is the scaling coefficient, *A*_*k*_ is the surface associated with vertex *k*, 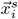 is the position of the sensor *i*, and 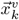 is the position of the vertex *k*. Note that we have not taken into account the dependence of the source-to-sensor decay on the orientation of the neuronal tissue, due to the lack of geometric information about the orientation in subcortical structures.

### State-space modeling

State-space modeling [41–43] forms the fundamental basis of dynamical systems theory [44, 45] and control engineering [46, 47], to describe and analyze system dynamics over time, capturing interactions within data and actively manipulating and regulating system behavior to steer the system towards desired states or trajectories [48, 49]. Nonlinear state-space modeling further enhances the modeling of complex systems, for instance to capture seizure onset, progression, and offset, by incorporating nonlinear relationships and dynamics [5, 9, 10, 50].

In this study, the state-space representation of the VEP model is given by a system of nonlinear differential equations as follows:

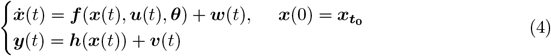

where 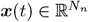 is the *N*_*n*_-dimensional vector of system states evolving over time, 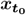 is the initial state vector at time *t* = 0, 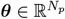 contains all the unknown evolution parameters, ***u***(*t*) stands for the external input, and 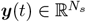 denotes the measured data subject to the measurement error ***v***(*t*). The process (dynamical) noise and the measurement noise denoted by ***w***(*t*) ∼ 𝒩 (0, σ^2^) and ***v***(*t*) ∼ 𝒩 (0, σ^*′*2^), respectively, are assumed to follow a Gaussian distribution with mean zero and variance σ and σ^*′*^, respectively. Moreover, ***f*** (.) is a vector function that describes the dynamical properties of the system i.e., summarizing the biophysical mechanisms underlying the temporal evolution of system states (here, govern by VEP model, Eq. 1) and ***h***(.) represents a measurement function i.e., the instantaneous mapping from system states to observations (here, the gain matrix, Eq. 2, and Eq. 3).

### Parameter estimation problem

In this study, it is assumed that the structure of the state-space model is known but that the associated model parameters θ ∈ {η_*i*_, *K*} with *I* ∈ {1, 2, …, *N*_*n*_} are unknown and need to be estimated from the available data. Although there are many strategies available for determining these parameters [10, 13, 14, 51–58], we focus on an approach based on global optimization. This method involves formulating an optimization problem that measures the difference between simulated data (produced by the model) and real observed data, aiming to adjust the model parameters for a better fit.

Given this context, to minimize the discrepancy between model predictions and observations, we employ the Root Mean Square Error (RMSE). A lower RMSE value indicates a better fit between model predictions and observed data, pointing to more accurate parameter estimation and enhanced model reliability.

Note that, due to the sparse placement of SEEG electrodes, the gain matrix is not of full rank (see S1 Fig), posing significant challenges for parameter estimation, particularly in accurately inferring the unknown combination of activity from neighboring brain regions near the sensor. These challenges include computational time, the reliability of estimating the epileptogenicity parameter, and the overall model inversion process. Consequently, the parameter estimation of dynamic models generally exhibits NP-hard complexity. Given the necessity to obtain a viable solution for the patient in a reasonable runtime, it is impractical to calculate for weeks to reach the optimal solution. Thus, in our global optimization approach, we have employed metaheuristics, local solvers, and HPC techniques, aiming to speedup the search as much as possible.

Subsequently, we can rearrange various configurations of the VEP model for benchmarking based on the connectivity matrix of a patient. Table 1 presents 6 different optimization problems and a summary of their main characteristics. As we are assuming no noise in problems 1 to 4, they are treated as ODEs, and their parameter estimation problems are deterministic. However, in the case of problems 5 and 6, we handle the noise, so the SDEs result in our cost function in the calibration being stochastic.

**Table 1.** Optimization problems for various configurations of the VEP model. The seizure propagation depends on the interplay between node dynamics (excitability) and network coupling (structure). The signals generated by VEP models on the brain region level are called source signals (see Eqs. 1). The measured signals from the electrodes are called sensor signals. To map the simulated sources from the brain regions to the sensors, the electromagnetic forward problem needs to be solved (see Eqs. 2 and 3). Due to the sparse placement of electrodes, the gain (lead-field matrix) used for mapping the source to the sensors is not of full rank.

| problem | cost function | description |
| --- | --- | --- |
| 1 | deterministic | Forward simulation of VEP model *with weak coupling* at *source level*, i.e., no propagation, and the gain matrix $G=I$. |
| 2 | deterministic | Forward simulation of VEP model *with weak coupling* at *sensor level*, i.e., no propagation, and a low rank gain matrix. |
| 3 | deterministic | Forward simulation of VEP model *with strong coupling* at *source level*, i.e., with propagation, and the gain matrix $G=I$. |
| 4 | deterministic | Forward simulation of VEP model *with strong coupling* at *sensor level*, i.e., with propagation, and a low rank gain matrix. |
| 5 | stochastic | Forward simulation of *SDE* VEP model with large $\tau$ (stiff equations) at *source level*, with propagation, and the gain matrix $G=I$. |
| 6 | stochastic | Forward simulation of *SDE* VEP model with large $\tau$ (stiff equations) at *sensor level*, with propagation, and a low rank gain matrix. |

### Cooperative optimization methods

The domain of High-Performance Computing (HPC) offers essential methodologies and advanced technologies designed to significantly boost the efficiency of classical optimization algorithms, particularly within the context of contemporary many-core infrastructures. These methods and techniques play a pivotal role in specific applications where computation speed is intrinsically linked to the patient’s well-being, underscoring the critical nature of rapid model calibration. In this context, messaging libraries like MPI emerge as key tools, providing an effective avenue for code parallelization. Concurrently, programming strategies such as the master-worker paradigm provide a streamlined and intuitive strategy for coordinating distributed optimization agents, further contributing to improve the overall efficacy and performance of numerical optimization methods.

Recently, leveraging the methodologies and technologies mentioned above, the Self-adaptive Cooperative enhanced Scatter Search (SaCeSS) [59] optimizer was presented as a competitive method to solve parameter estimation problems in large-scale nonlinear dynamic models. SaCeSS is a cooperative parallel (using an island-based model) implementation of an evolutionay algorithm, enhanced scatter search (eSS) [60, 61]. Independent eSS instances (workers) run in parallel in a cooperative fashion, and a master process manages communications between them. Each parallel eSS instance is a hybrid method, combining global search with calls to an efficient local search (LS) method (in this study, Dynamic Hill Climbing, DHC [62]).

During the execution of SaCeSS, if a worker explores a solution that improves the best-known one (considering all workers), this solution is asynchronously sent to the master, assessing its quality and determining whether it should be propagated to other MPI processes. This hybrid optimization strategy is enriched with several innovative mechanisms. These include: (i) the establishment of asynchronous collaboration amongst parallel processes, ensuring a seamless and efficient exchange of information; (ii) the integration of both coarse and fine-grained parallelism, providing a balanced and versatile computational structure; and (iii) the implementation of self-tuning strategies, which autonomously improve performance and adapt to varying conditions, ultimately enhancing the reliability and efficiency of the search process.

To provide a visual representation of the communication and adaptation mechanisms in SaCeSS, Figure 1 offers a detailed pictorial overview. In previous research, SaCeSS demonstrated good performance and robustness in addressing complex, large-scale model calibration problems within the domain of computational systems biology [63]. Leveraging these strengths, here we have tailored SaCeSS to tackle the specific optimization challenges associated with parameter estimation in VEP models.

**Fig. 1.**
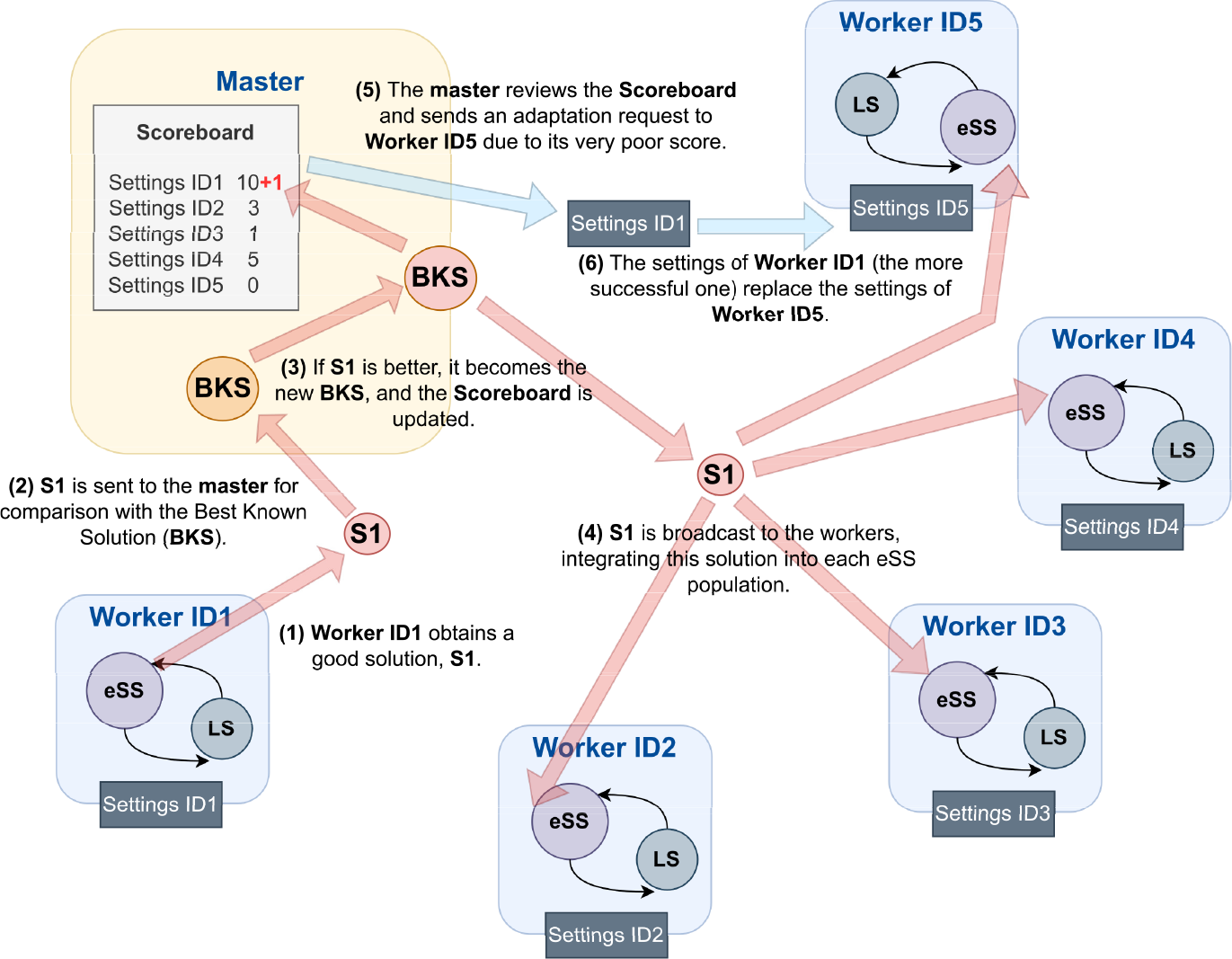
General Overview of SaCeSS. This method employs a parallel cooperative scheme based on a master-worker strategy. An example of solution propagation flow is illustrated: when a worker obtains a good solution, it is shared with the rest of the workers through the master. Additionally, the master implements an adaptation mechanism to replace the settings of those workers with poor performance (as registered in a scoreboard), helping to improve their performance.

### Scope and contributions

In this study, we employed SaCeSS algorithm to calibrate various configurations of the VEP model, thereby demonstrating the potential of this solver for rapid parameter estimation in such contexts. As the VEP models serve as personalized brain network models tailored to individual patients, they can lead to more informed clinical decision-making and optimize surgical strategies. For this reason, the primary objective is to ascertain whether our proposed approach can address the associated optimization problem within a reasonable computational time to avoid excessive waiting times for the patient.

We also evaluate the parallel efficiency and scalability of SaCeSS in the context of VEP models using two types of HPC infrastructures, a supercomputer and a PC workstation. Based on these results, we analyze the computational resources required to obtain fast and sufficiently accurate estimations, which is important to prevent extended delays in delivering medical results in a real-world application. Further, we also compare SaCeSS with other competitive parallel optimizers.

Additionally, since the calibration of large-scale models always contains some degree of uncertainty due to non-identifiabilities, we extend SaCeSS with a method of uncertainty quantification based on a recent ensemble approach [64, 65].

Finally, while SaCeSS is equipped with a self-adjusting mechanism for its settings, there remain certain global hyperparameters associated with its overall cooperation and adaptation functionalities. To enhance SaCeSS’s robustness and efficiency even further, here we have used Optuna, a hyperparameter tuning tool [66]. This tool leverages Bayesian optimization techniques to determine the most effective global hyperparameters considering the specific nature of the VEP parameter estimation challenges.

## Results

In this section, our aim is to assess the performance of our proposed parameter estimation method in VEP models. To achieve this, we tested the SaCeSS method with the different benchmarks presented in Table 1. For each optimization problem, SaCeSS was run using the same computational resources: 12 parallel processors within a specific time threshold. Due to the non-deterministic nature of SaCeSS, it was necessary to perform multiple runs (10 times per problem) to ensure a comprehensive assessment. Recognizing the differing complexities of each benchmark and aiming to provide adequate time for convergence, we established the following stopping times (wall time limits) in the FT3 supercomputer: 3 hours for problem 1, 18 hours for problems 2-4, and 39 hours for problems 5 and 6.

Figs 2, 3, 4 present the VEP configurations and estimations for different spatial maps of epileptogenicity, at source and sensors levels (see Table 1). For each problem, we have shown the whole-brain simulations (see Eq. 1) at source level and corresponding SEEG simulations at sensor level (see Eq. 2). Moreover, the trajectories in the phase-plane for different regions are illustrated. These results aim to showcase the capabilities of SaCeSS in capturing the true mechanism underlying seizure initiation and propagation from a dynamical systems theory perspective. We have utilized the confusion matrix, which was computed based on the inferred excitability η_*i*_, to report the estimation accuracy for three node types: Epileptogenic Zone (EZ), Propagation Zone (PZ), and Healthy Zone (HZ).

**Fig. 2.**
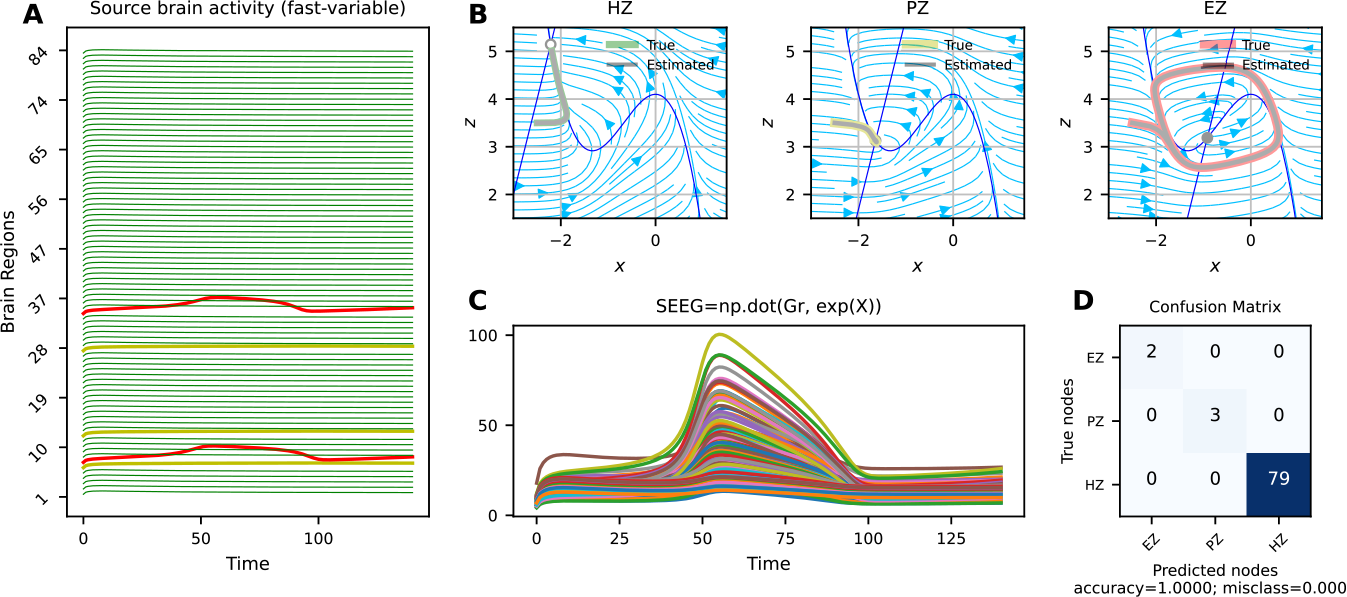
Best solutions obtained by SaCeSS using 12 processors for the deterministic VEP model with weak coupling at source and sensor levels (problems 1 and 2). (**A**) The simulated fast activities across brain regions at source level used for optimization. No propagation is observed due to weak coupling. (**B**) The true and estimated trajectories at phase-plane for three node types as Epileptogenic Zone (EZ, in red), Propagation Zone (PZ, in yellow), and Healthy Zone (HZ, in green). (**C**) The envelope of the simulated SEEG signals at the sensor level. (**D**) The confusion matrix indicates 100% accuracy in the estimation of three node types (EZ/PZ/HZ) based on SEEG signals.

**Fig. 3.**
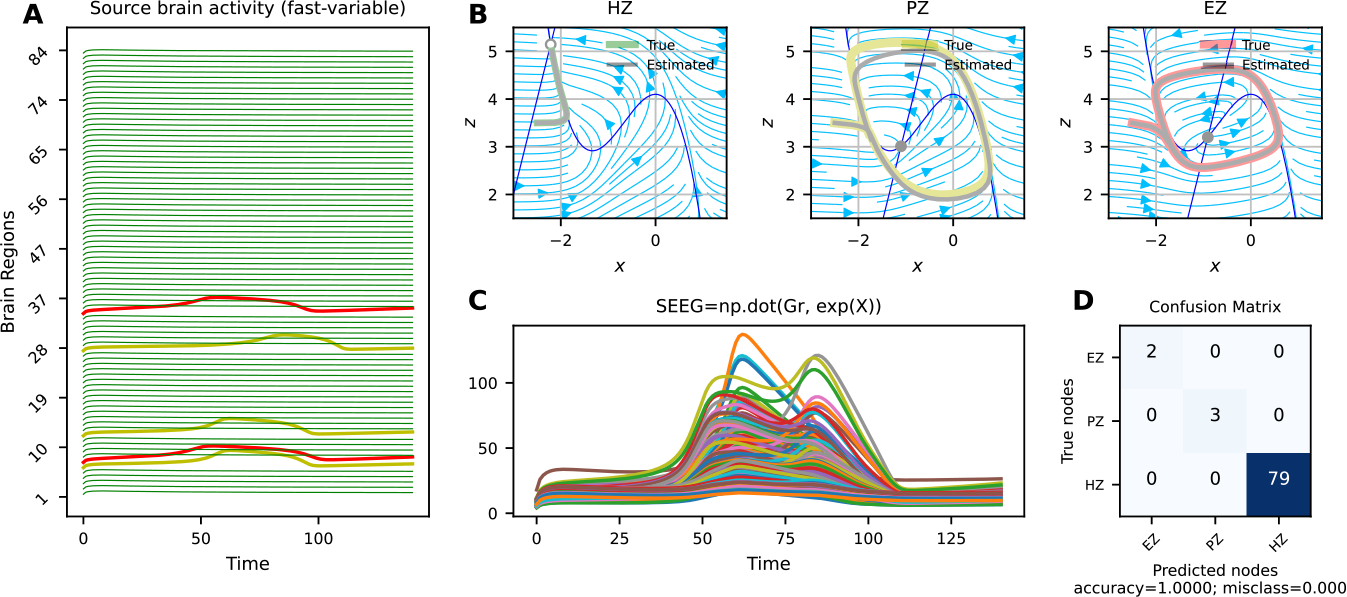
Best solutions obtained by SaCeSS using 12 processors for deterministic VEP model with strong coupling at source and sensor levels (problems 3 and 4). **A**) The simulated fast activities across brain regions at source level used for optimization. Due to strong coupling, the seizure propagates from epileptogenic zone (in red) to other brain regions (in yellow). (**B**) The phase-plane displayed both the actual and predicted trajectories for three node categories as EZ, PZ, and HZ. (**C**) The simulated SEEG signals at the sensor level. (**D**) The confusion matrix that illustrates 100% accuracy for the estimation based on SEEG signals.

**Fig. 4.**
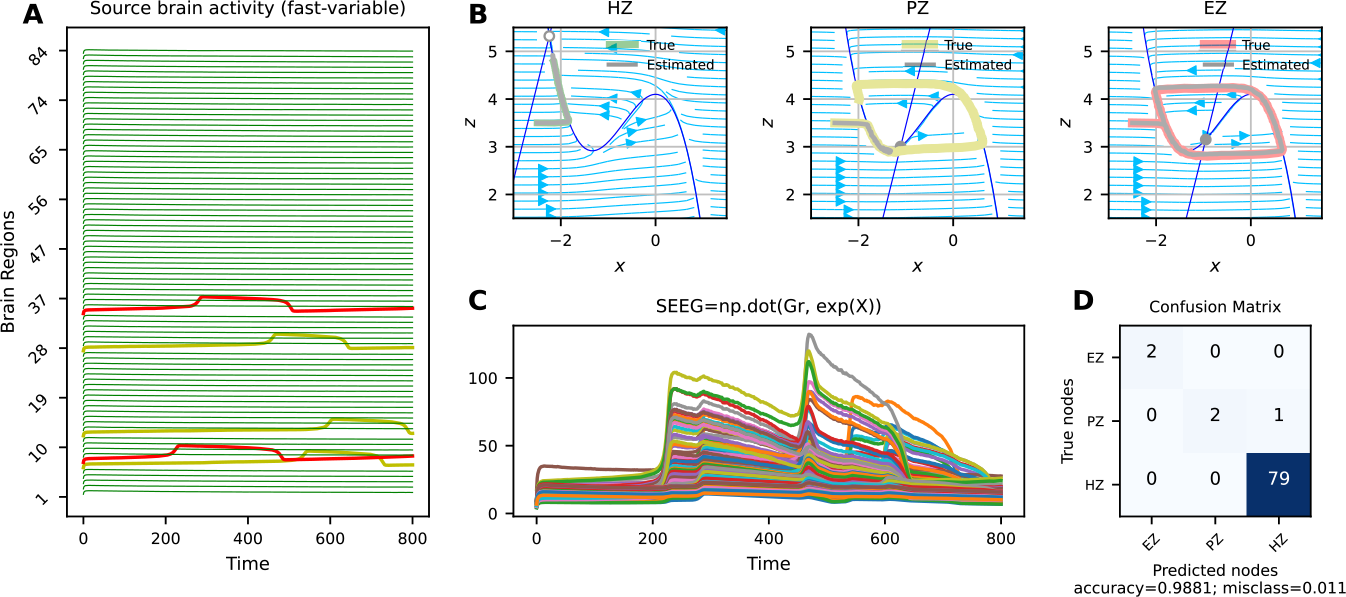
Best solutions obtained by SaCeSS using 12 processors for stochastic VEP model with stiff equations at source and sensor levels (problems 5 and 6). **A**) The simulated fast activities across brain regions with large sizure lenght (due to large time scale separation τ) at source level used for optimization. Due to strong coupling, the seizure propagates from EZ (in red) to PZ (in yellow). (**B**) The true and estimated trajectories in phase-plane for EZ, PZ, and HZ. (**C**) The simulated SEEG signals at the sensor level. (**D**) The confusion matrix that illustrates that one of the PZ is mis-classified as HZ.

Fig 2 shows the estimation result for the deterministic VEP model with weak coupling, i.e., no seizure recruitment from EZ to PZ. Fig 2**A** displays the simulated fast activities across brain regions at source level (cf., *x*_*i*_ variables in Eq. 1). In this scenario, two regions with high excitability η_*i*_ are identified as EZ (shown in red), while three regions are designated as candidate regions with excitability close to the critical value for seizure propagation (PZ, shown in yellow). The remaining regions with low excitability are classified as healthy (HZ, shown in green). Fig 2**B** displays the true and estimated trajectories in the phase-plane. In the phase-planes depicted, the x- and z-nullclines are highlighted in blue, and the intersection of these nullclines indicates the fixed point of the system. A full circle represents a stable fixed point, while an empty circle represents an unstable fixed point. From this figure, we can observe that the trajectory of an HZ is attracted to the stable fixed point of the system (located on the left branch of the cubic x-nullcline), indicating that it does not trigger an epileptic seizure. For a PZ, the coupling and the value of excitability, which is close to the critical value of epileptogenicity, cause the z-nullcline to move downward. However, the coupling is not sufficient for propagation as the system’s fixed point remains stable. For the EZ, the system exhibits an unstable fixed point due to the high value of excitability. In this regime, Epileptor possesses a limit cycle and the seizure triggers autonomously. We can observe that in all cases, the estimated trajectories closely follow the true trajectories due to the accurate estimation of excitability (see Table 2). Fig 2**C** shows the simulated SEEG signals at the sensor level given a low-rank sparse gain matrix. Fig 2**D** indicates a 100% accuracy in the estimation of three node types (EZ/PZ/HZ) based on SEEG signals. Overall, these results indicate that SaCeSS is able to accurately capture the system dynamics for different node types in the absence of noise in the VEP model.

**Table 2.** Results for different parallel GO methods considering the set of VEP problems. Each solver was executed 10 times (different parallel jobs), with different stopping times depending on the problem. Resulted obtained using 12 parallel processors in the FT3 supercomputer.

| problem | stopping time | best fx | mean fx ± std | problem | stopping time | best fx | mean fx ± std |
| --- | --- | --- | --- | --- | --- | --- | --- |
| **SaCeSS** |  |  |  | **PS-CMA-ES** |  |  |  |
| 1 | 3 h | 5.30E-07 | 5.72E-07 ± 2.56E-08 | 1 | 3 h | 1.70E-13 | 1.76E-13 ± 4.99E-15 |
| 2 | 18 h | 2.20E-03 | 2.71E-03 ± 4.30E-04 | 2 | 18 h | 1.70E+00 | 6.38E+00 ± 6.49E+00 |
| 3 | 18 h | 4.93E-07 | 5.21E-07 ± 1.75E-08 | 3 | 18 h | 1.59E-13 | 1.70E-13 ± 5.55E-15 |
| 4 | 18 h | 3.39E-03 | 1.01E-02 ± 4.82E-03 | 4 | 18 h | 2.82E+00 | 7.28E+00 ± 3.88E+00 |
| 5 | 39 h | 7.35 | 7.74 ± 3.09E-01 | 5 | 39 h | 22.48 | 24.46 ± 1.19 |
| 6 | 39 h | 632.38 | 719.24 ± 130.47 | 6 | 39 h | 1143.70 | 1302.89 ± 171.17 |
| **asynPDE** |  |  |  | **SaCeSS optuna** |  |  |  |
| 1 | 3 h | 2.15E-05 | 8.68E-05 ± 1.01E-04 | 1 | 3 h | 5.00E-07 | 5.47E-07 ± 3.31E-08 |
| 2 | 18 h | 1.11E+00 | 1.35E+01 ± 1.37E+01 | 2 | 18 h | 3.31E-04 | 7.32E-04 ± 3.15E-04 |
| 3 | 18 h | 4.31E-13 | 5.99E-05 ± 9.18E-05 | 3 | 18 h | 4.99E-07 | 5.15E-07 ± 1.17E-08 |
| 4 | 18 h | 1.07 | 19.60 ± 9.47 | 4 | 18 h | 4.90E-04 | 8.84E-04 ± 3.75E-04 |
| 5 | 39 h | 27.01 | 30.95 ± 3.32 | 5 | 39 h | 7.22 | 7.69 ± 0.25 |
| 6 | 39 h | 961.30 | 1195.68 ± 109.48 | 6 | 39 h | 535.88 | 792.44 ± 197.92 |

Fig 3 shows the estimation result for the deterministic VEP model but with strong coupling, i.e., with seizure recruitment from EZ to PZ. Fig 3**A** displays the simulated fast activities across brain regions at the source level, where the seizure propagates to the candidate brain regions as PZ. Fig 3**B** The seizure recruitment to these regions is due to the network effects, as their steady-state equilibrium is close to the bifurcation value. Here, due to the sufficient coupling strength and the value of excitability which is close to the critical value of epileptogenicity, the z-nullcline moves down, causing a bifurcation thereby allowing the seizure to propagate. This indicates that seizure propagation depends on the interplay between the brain region’s excitability (node dynamics), and the network coupling (parameter K). Fig 3**C** and Fig 3**D** show the observed SEEG signals and the accuracy of the estimation. These results indicate that SaCeSS is able to accurately capture the system dynamics for different node types with strong coupling in the VEP model.

Fig 4 shows the estimation result for the stochastic and stiff VEP model, i.e., with a long seizure envelope (see Fig 3**A**) due to a large time scale separation (parameter τ in Eq. 1). In particular, we have considered a high value of noise dynamics, which is a zero-mean Gaussian noise with a standard deviation of 0.1. Therefore, given a known values of excitability and coupling, the seizure propagation is random and depends on the brain state dependency caused by the noise dynamics. Fig 3**B** shows that the system dynamics of regions corresponding to EZ and HZ are accurately estimated. However, the estimated trajectory corresponding to one of PZs dampens to a stable fixed point, despite the presence of a limit cycle in the observation. Fig 3**C** shows the simulated SEEG signals used for optimization, where one of the PZ regions is misclassified as HZ, as indicated by the confusion matrix (Fig 3**D**). This result indicates that although SaCeSS can handle the very fast-changing components in the VEP model, accurately estimating the noise can be challenging due to the use of an error function such as RMSE for optimization.

Table 2 lists the best costs (RMSE) for each problem, alongside average and deviation values, where problems from 1 to 4 (ODEs) approached zero. In contrast, the convergence of problems 5 and 6 (SDEs) is not close to zero as for the models based on ODEs, but the estimations are close to ground true values.

We further assessed the scalability of SaCeSS on two platforms: the FinisTerrae III (FT3) supercomputer (details at https://www.cesga.es/en/), and a DELL Precision 5820 workstation equipped with 18 cores (Intel i9-10980XE at 3.00GHz). We conducted ten runs of each optimization problem using both the sequential eSS and the parallel SaCeSS solvers, employing 6, 12, and 24 processors—although only 6 and 12 cores were used in the case of the DELL Precision 5820. This exercise served to highlight the performance advantages of SaCeSS over eSS and offered valuable observations regarding the behavior of SaCeSS with varying numbers of parallel processors (workers).

The detailed results of this analysis are shown in Table 3, where, in general, all SaCeSS configurations achieved fitness functions close to zero, except for problems 5 and 6, where the RMSE metric is unable to capture the noisy nature of the observation at the sensor level. The results indicate that increasing the number of processors (workers) leads to an enhancement in the accuracy of solutions. This improvement is also mirrored in the reduction of mean and deviation values. Figure 5 provides additional details: the boxplots show the spread of the best solutions obtained, and the convergence curves (evolution of objective function versus time considering the best run of each method) illustrate the speedup gains with more processors. The bootstrapping on different runs complements this observation, indicating that the distribution of solutions has reduced its variability and is pushed towards better results.

**Table 3.**
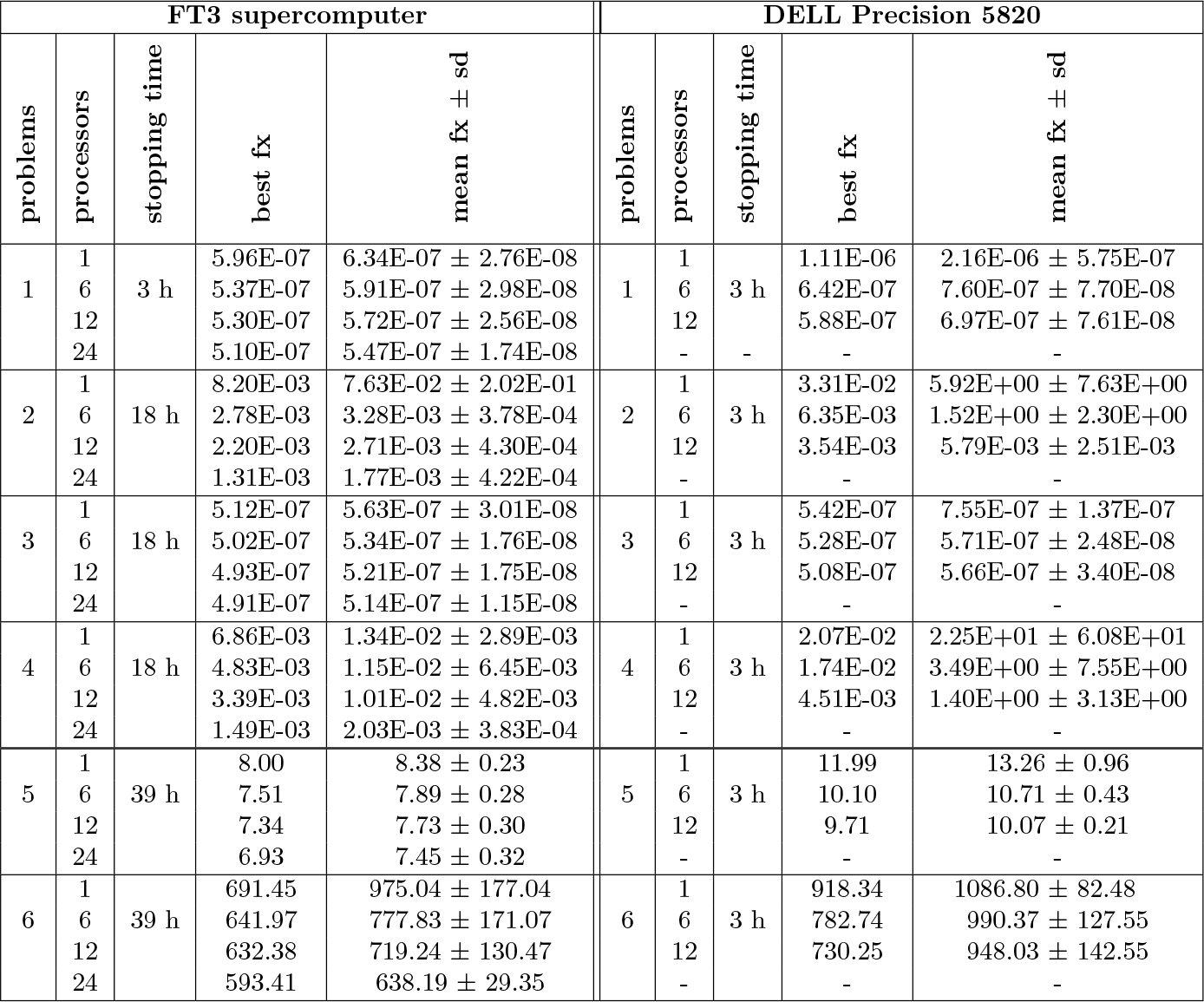
Scalability of SaCeSS in an supercomputing infrastructure (FT3) and in a desktop workstation (DELL Precision 5820). SaCeSS was executed 10 times using different number of processors, varying the stopping time for each problem. In the case of one processor, the sequential enhanced scatter search (eSS) solver was used.

**Fig. 5.**
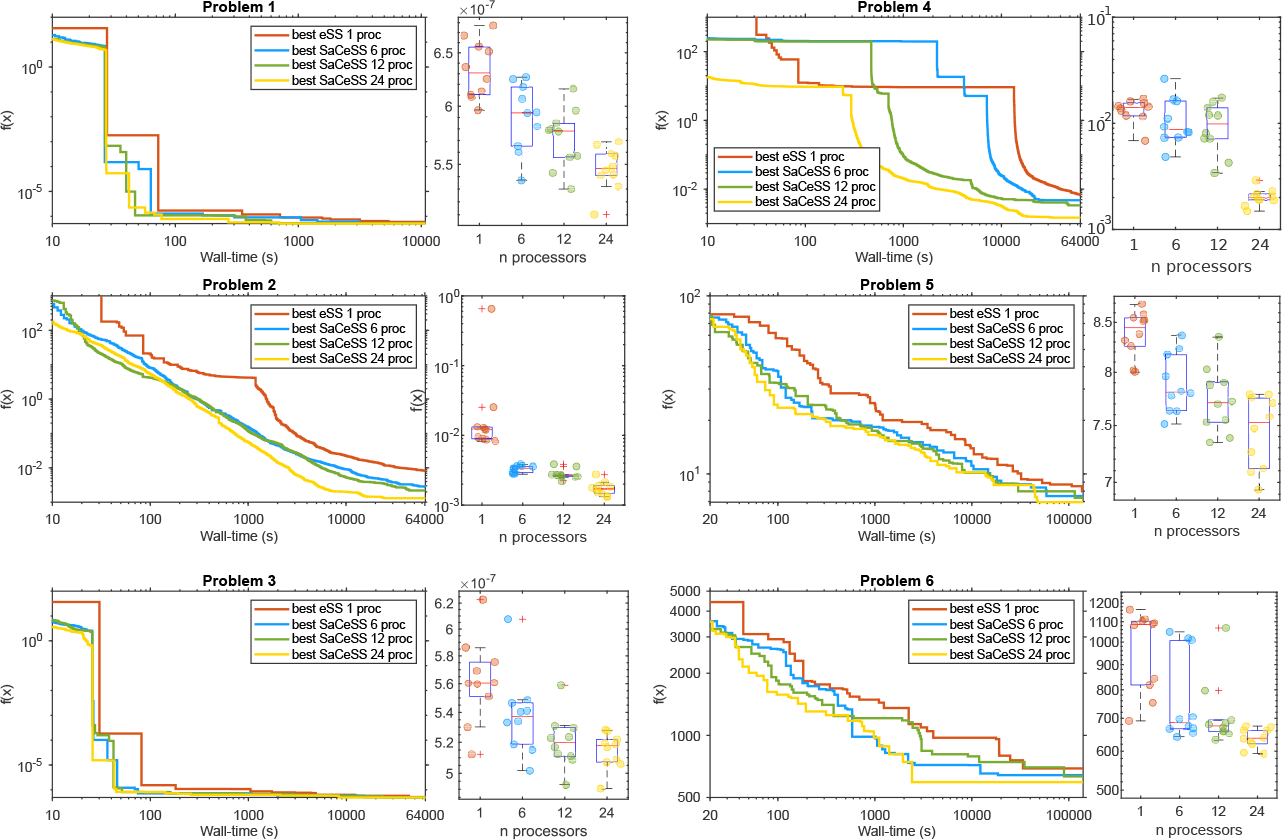
Scalability analysis of SaCeSS on the Finisterrae III supercomputer. Each convergence curve represents the best run for the sequential method (eSS) and for SaCeSS using 6, 12, and 24 processors. The boxplots illustrate the spread in the solutions obtained in repeated runs (using different number of parallel workers for the case of SaCeSS).

Additionally, we benchmarked SaCeSS against two competitive parallel global optimizers which have previously shown good performance in various computational biology problems: the Particle Swarm CMA Evolution Strategy (PS-CMA-ES) [67] and the asynchronous parallel Differential Evolution (asynPDE) [68]. Maintaining the same computational testbed with 12 parallel processors per run and the previously mentioned time thresholds, we analyzed for these solvers the quality of the best solutions and the associated means and deviations. Table 2 presents a comprehensive summary of results, and Figure 6 shows the convergence curves and final solution spread (with boxplots). These reults unequivocally demonstrate the enhanced performance of our proposed method over both asynPDE and PS-CMA-ES in various aspects of solving VEP problems. Our method exhibits a more rapid convergence rate, better robustness in the estimations (as indicated by a lower dispersion in results across multiple runs), and the ability to find superior solutions. The sole exceptions are observed when addressing the deterministic VEP at the source level (problems 1 and 3), where the estimations are much easier due to the full identifiability of all parameters involved. In these particular problems, PS-CMA-ES yields results very close to zero (in the order of 1*E* − 13), outperforming SaCeSS, which provides results in the order of 1*E* − 7. While SaCeSS converges more swiftly, it tends to level off at these values. Nonetheless, it is important to note that these fits are exceptionally accurate and, from a practical standpoint, they are virtually indistinguishable.

**Fig. 6.**
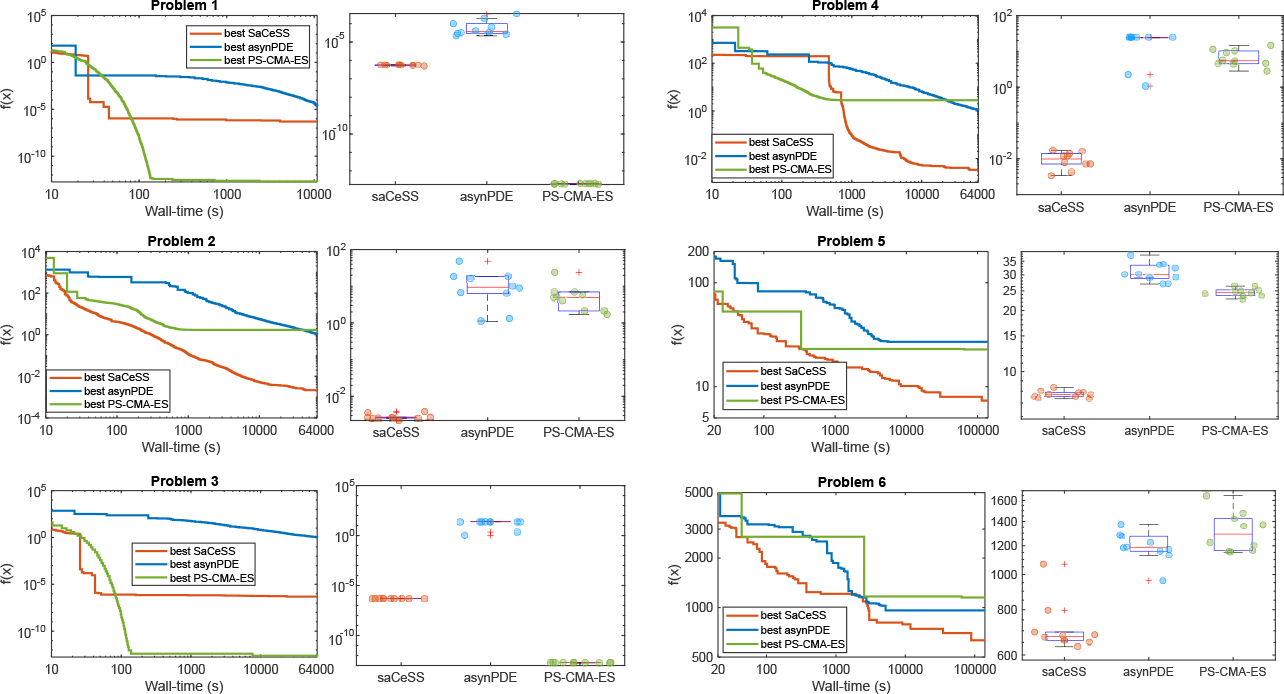
Comparison of SaCeSS with other parallel optimizers. Each convergence curve represents the best run for SaCeSS, asynPDE, and PS-CMA-ES, all using 12 processors. The boxplots illustrate the dispersion in the solutions obtained with the aforementioned solvers. Each colored spot within the boxplots represents the solution cost obtained by a single run for each solver.

We also explored the us of a Bayesian optimization framework (via Optuna [66]) to further improve the performance of SaCeSS in the VEP benchmarks. The objective was to fine-tune specific configuration settings related to the cooperation mechanism. This approach requiered the formulation of a new mixed-integer optimization problem with six decision variables associated with the configuration options to be tuned. The objective was to minimize a cost function defined by the geometric mean of cost values from five independent SaCeSS runs. Each of these five runs used 12 processors with stopping criteria only based on time (30 seconds). We used this threshold because it is the time required to reach quality solutions in Problem 1. In other words, we used the Bayesian optimization scheme provided by Optuna to obtain the best SaCeSS hyperparameters for Problem 1. This process required 13 hours to complete and was repeated 10 times due to the stochastic nature of Bayesian optimization. We only performed this Bayesian tuning for Problem 1, the easiest case study. The computational cost of repeating this fine-tuning for the other problems would be very significant.

For this reason, we applied the best settings obtained during the calibration of problem 1 to the others. assuming that the Bayesian fine-tuning would generalize well. The results are reported in Table 2, where the fine-tuned SaCeSS outperformed the original SaCeSS configuration in terms of both the best solution and its mean. Figure 7 provides a side-by-side comparison, visually illustrating the differences in solution spread and convergence curves (across all runs) between the standard and the fine-tuned SaCeSS. The latter converged faster and to better solutions in problems 1-4, reduced the spread in the results. However, the results for problems 5 and 6 were not statistically different, indicating that the SaCeSS default self-adaptive mechanisms are capable of handling the more challenging problems.

**Fig. 7.**
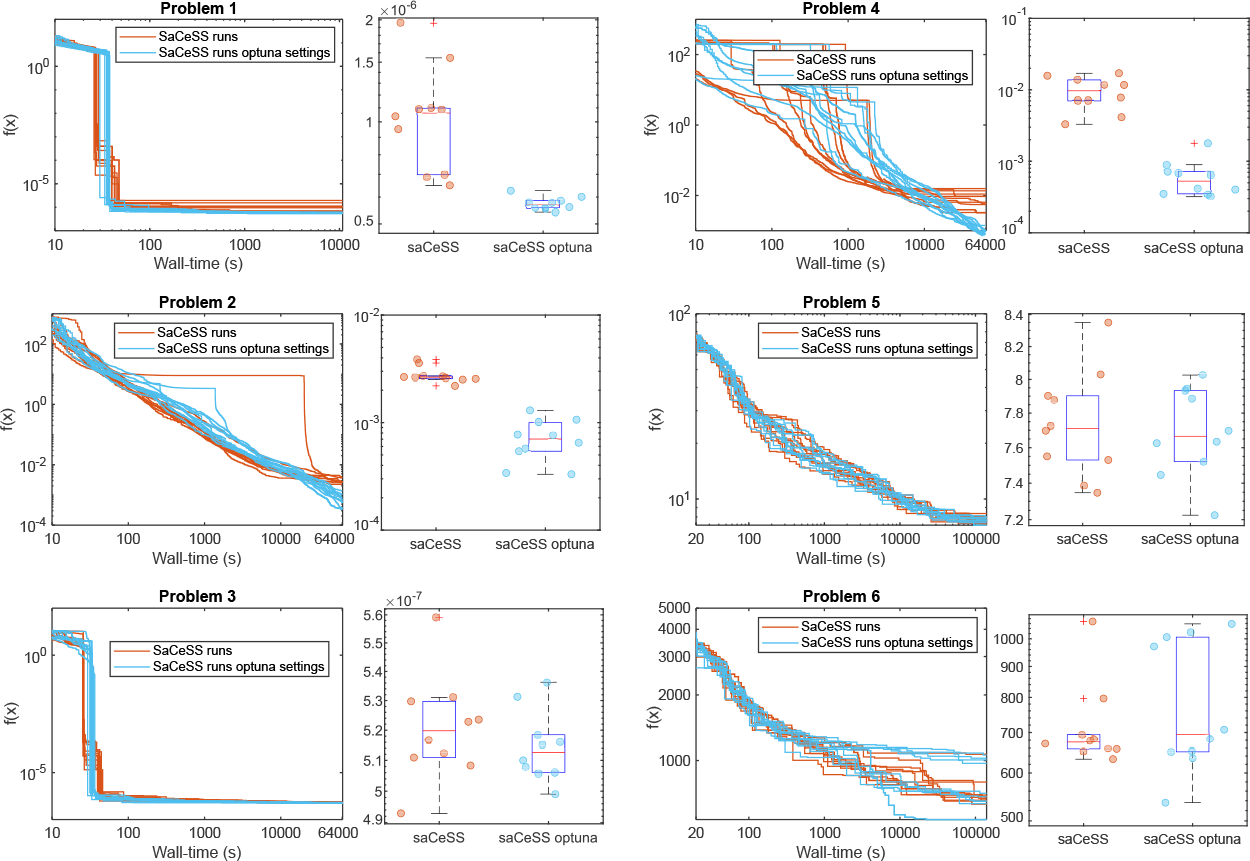
Comparison between the original and the fine-tuned SaCeSS. Convergence curves for original (orange) and fine-tuned SaCeSS (blue), plus boxplots illustrating the solutions spread.

Regarding uncertainty quantification, we used an approach based on ensemble modeling [64, 65]. This method takes samples from the SaCeSS optimization in the vicinity of the global solution, building an ensemble of calibrations which represent equally well the data. During the optimizations, SaCeSS stores all the explored solutions of certain quality (defined by a threshold). After convergence, our workflow selects a representative subsample of the stored solutions, normalizes it, and then illustrate the variability in the estimated parameters for each problem using parallel coordinate graphs. These results are shown in Figure 8, wherein the y-axis, the time-varying distribution percentiles are shown as shaded red bands around a central black line (the median). The x-axis represents the parameters’ index in the VEP models: the level of epileptogenicity η_*i*_ at each brain region and the global coupling *K* (for the label names and indices of the sub-divided brain regions, see Table 2). Due to the sparsity of the gain matrix, we observe a higher level of uncertainty in the estimation at the sensor level (problems 2, 4, and 6) compared to the source level (problems 1, 3, and 5).

**Fig. 8.**
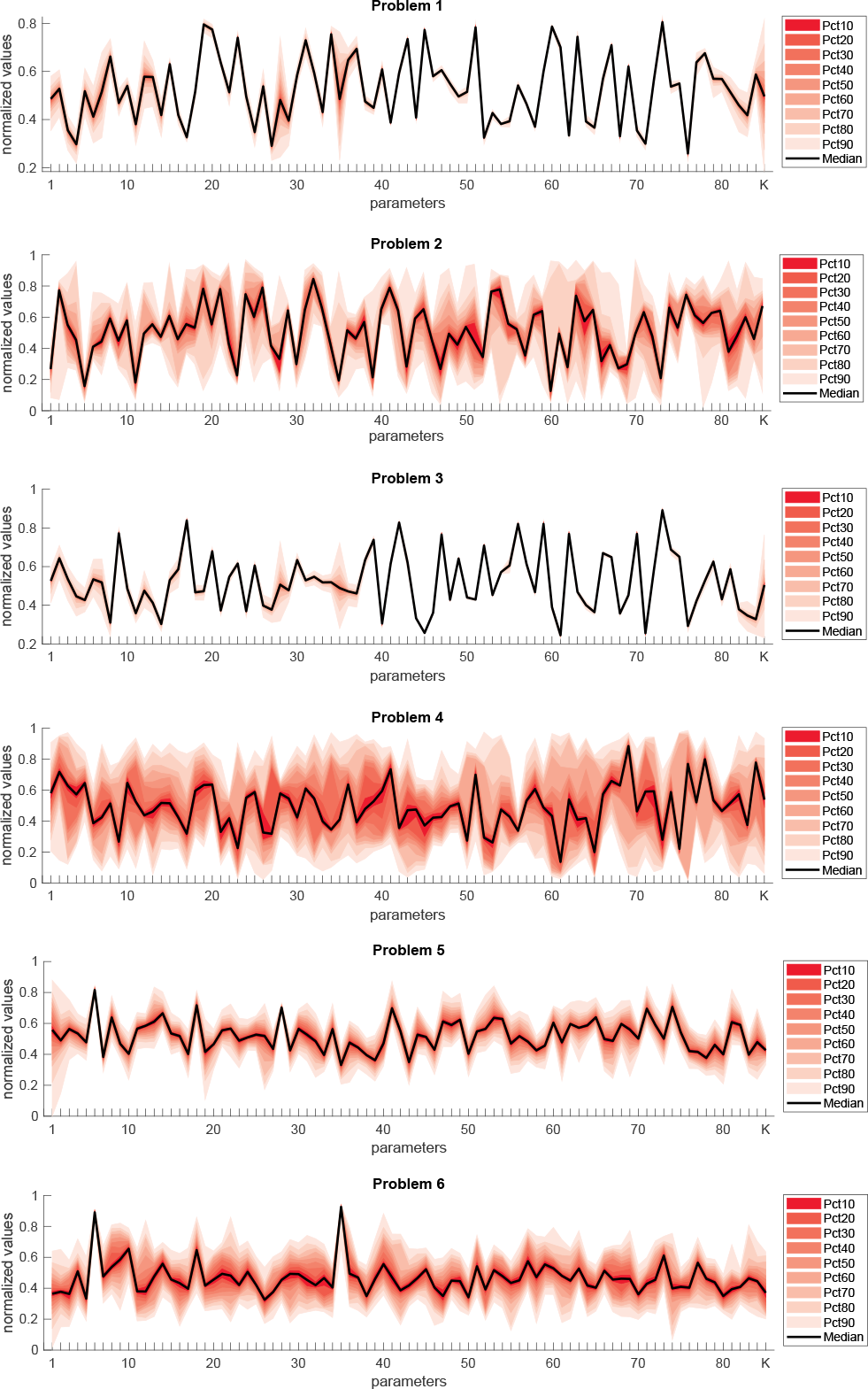
Uncertainty quantification of the parameters estimated using SaCeSS: dispersion as parallel coordinates plots. On the y-axis, the time-varying distribution percentiles are manifested as shaded red bands encircling a central black line, which signifies the median. The x-axis represents the parameters in the VEP models: the level of epileptogenicity η_*i*_ at each brain region and the global coupling *K*.

## Discussion

The accurate parameter estimations in different configurations of the VEP model, as shown in Table 2 and Figures 2, 3 and 4 highlight the capability of SaCeSS to solve high-dimensional problems within a reasonable computational time, using just 12 processors (additional details can be found at https://doi.org/10.5281/zenodo.10057789). The convergence curves further substantiate this observation: in different deterministic problems, SaCeSS exhibit fast convergence while obtaining solutions close to the global optimum. However, in stochastic problems, SaCeSS presents a less efficient behavior due to the complexity introduced by dynamical noise. In particular, the solution for Problem 6 did not estimate the phase-plane trajectories in the PZ region adequately. Thus, in order to enhance efficiency when addressing stochastic models, a promising strategy could be to study the integration of SaCeSS with Bayesian optimization methods, known for their proficiency in such challenges, at least for problems of moderate number of parameters.

Moreover, our method demonstrates versatility across different computational infrastructures. The original SaCeSS was designed for high-performance computing environments because of its parallel nature and scalability, i.e. adding parallel workers to the search increases the probability of finding a better solution faster. Nevertheless, SaCeSS also delivers steadfast performance on smaller-scale systems, such as desktop workstations equipped with a moderate quantity of cores and memory, even when operating under stringent time limitations (see Table 3, “DELL Precision 5820” results). This flexibility ensures accessibility for a broad user base, including those with limited experience in high-performance computing, allowing them to fully exploit the advantages offered by our methodology.

In the comparison of SaCeSS with two other leading parallel solvers (asynPDE and PS-CMA-ES), as showcased in Table 2, our approach demonstrates a consistent advantage over these competitors across multiple criteria in addressing VEP problems. These results highlight the competitive edge of cooperative parallel methods like SaCeSS against other strategies utilizing more conventional parallelization techniques. Moreover, considering the scarcity of parallel global optimization methods, and the complexities entailed in handling estimation problems involving differential equations, this comparative analysis emerges as a significant and informative contribution in itself.

Finally, the fine-tuning of SaCeSS using Bayesian optimization has resulted in significant improvements, particularly in the VEP problems with a deterministic cost function. We regard this as a first step exploring a promising field: the fine-tuning of cooperative parallel methods for challenging global optimization problems. In this particular context, as opposed to sequential algorithms, the settings to be fine-tuned are related to “social” features of multi-agent systems, such as communications among different workers. Therefore, several open questions remain for future consideration, including the calibration of stochastic problems given their high computational cost, and enhancing this procedure through a sensitivity analysis to determine the most impactful settings for inclusion or exclusion during fine-tuning.

In summary, driven by the challenge of estimating parameters in a series of progressively complex whole-brain network models related to epilepsy, this study started with the objective of selecting a resilient and efficient global optimization solver. As a second objective, we wanted a solver capable of leveraging parallel computing across various platforms, ranging from personal desktop workstations to supercomputers.

Our investigation revealed the robustness and efficiency of the SaCeSS parallel solver in addressing the complex parameter estimation problems associated with VEP models. This algorithm outperformed other parallel solvers in most VEP benchmark problems and exhibited enhanced performance when scaling the number of parallel processors. Additionally, using a Bayesian framework for hyperparameter tuning, we were able to further improve its performance. We also extended SaCeSS with a mechanism to store and post-process the parameter space sampled in the vicinity of the global solution and, after convergence, use the sampling to perform approximate uncertainty quantification. This extension allowed a computationally efficient assessment of the variability in the estimated parameters. Overall, these findings collectively highlight the potential of our global optimization approach in contributing to more informed clinical decision-making through fast and accurate parameter estimation in VEP models.

### Information Sharing Statement

Detailed results and the main source codes needed to reproduce them are available on Zenodo https://doi.org/10.5281/zenodo.10057789. For the most recent version of the code, visit GitHub at https://github.com/ins-amu/BVEP/tree/master/Optimization.

## Acknowledgments

The authors acknowledge CESGA (Centro de Supercomputación de Galicia) for providing access to its FinisTerrae III supercomputer. J.R.B. and D.R.P. acknowledge financial support from grant PID2020-117271RB-C22 (BIODYNAMICS) funded by MCIN/AEI/10.13039/501100011033. M.H and V.J acknowledge the funding by the French National Research Agency (ANR) as part of the second “Investissements d’Avenir” program, ANR-17-RHUS-0004, EPINOV (https://anr.fr), and Human Brain Project SGA3 (https://ec.europa.eu/programmes/horizon2020).

## Supporting information

**S1 Fig.**
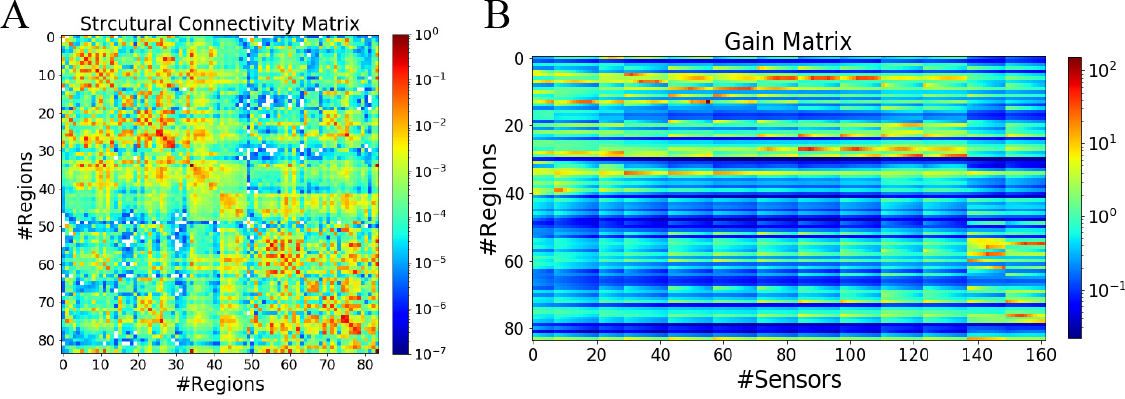
The structural connectivity and gain matrix used in VEP model. (**A**) An exemplary of structural connectivity (SC) matrix, whose entries represent the connection strength between the brain regions, is derived from diffusion MRI tractography. Using Desikan-Killiany parcellation, the brain of the patient is parcellated into 84 different regions comprising 68 cortical regions and 16 subcortical structures. (**B**) Rectangular gain (lead-field or projection) matrix of a randomly selected patient with partial epilepsy. Each element represents the inverse-squared distance between the region and the sensor. By virtue of sparse implantation, most elements of gain matrix are close to zeros.

**S2 Table.** Labels and indices of sub-divided brain regions.

| # | label | # | label |
| --- | --- | --- | --- |
| 01 | ctx-lh-bankssts | 43 | Right-Thalamus-Proper |
| 02 | ctx-lh-caudalanteriorcingulate | 44 | Right-Caudate |
| 03 | ctx-lh-caudalmiddlefrontal | 45 | Right-Putamen |
| 04 | ctx-lh-cuneus | 46 | Right-Pallidum |
| 05 | ctx-lh-entorhinal | 47 | Right-Hippocampus |
| 06 | ctx-lh-fusiform | 48 | Right-Amygdala |
| 07 | ctx-lh-inferiorparietal | 49 | Right-Accumbens-area |
| 08 | ctx-lh-inferiortemporal | 50 | ctx-rh-bankssts |
| 09 | ctx-lh-isthmuscingulate | 51 | ctx-rh-caudalanteriorcingulate |
| 10 | ctx-lh-lateraloccipital | 52 | ctx-rh-caudalmiddlefrontal |
| 11 | ctx-lh-lateralorbitofrontal | 53 | ctx-rh-cuneus |
| 12 | ctx-lh-lingual | 54 | ctx-rh-entorhinal |
| 13 | ctx-lh-medialorbitofrontal | 55 | ctx-rh-fusiform |
| 14 | ctx-lh-middletemporal | 56 | ctx-rh-inferiorparietal |
| 15 | ctx-lh-parahippocampal | 57 | ctx-rh-inferiortemporal |
| 16 | ctx-lh-paracentral | 58 | ctx-rh-isthmuscingulate |
| 17 | ctx-lh-parsopercularis | 59 | ctx-rh-lateraloccipital |
| 18 | ctx-lh-parsorbitalis | 60 | ctx-rh-lateralorbitofrontal |
| 19 | ctx-lh-parstriangularis | 61 | ctx-rh-lingual |
| 20 | ctx-lh-pericalcarine | 62 | ctx-rh-medialorbitofrontal |
| 21 | ctx-lh-postcentral | 63 | ctx-rh-middletemporal |
| 22 | ctx-lh-posteriorcingulate | 64 | ctx-rh-parahippocampal |
| 23 | ctx-lh-precentral | 65 | ctx-rh-paracentral |
| 24 | ctx-lh-precuneus | 66 | ctx-rh-parsopercularis |
| 25 | ctx-lh-rostralanteriorcingulate | 67 | ctx-rh-parsorbitalis |
| 26 | ctx-lh-rostralmiddlefrontal | 68 | ctx-rh-parstriangularis |
| 27 | ctx-lh-superiorfrontal | 69 | ctx-rh-pericalcarine |
| 28 | ctx-lh-superiorparietal | 70 | ctx-rh-postcentral |
| 29 | ctx-lh-superiortemporal | 71 | ctx-rh-posteriorcingulate |
| 30 | ctx-lh-supramarginal | 72 | ctx-rh-precentral |
| 31 | ctx-lh-frontalpole | 73 | ctx-rh-precuneus |
| 32 | ctx-lh-temporalpole | 74 | ctx-rh-rostralanteriorcingulate |
| 33 | ctx-lh-transversetemporal | 75 | ctx-rh-rostralmiddlefrontal |
| 34 | ctx-lh-insula | 76 | ctx-rh-superiorfrontal |
| 35 | Left-Cerebellum-Cortex | 77 | ctx-rh-superiorparietal |
| 36 | Left-Thalamus-Proper | 78 | ctx-rh-superiortemporal |
| 37 | Left-Caudate | 79 | ctx-rh-supramarginal |
| 38 | Left-Putamen | 80 | ctx-rh-frontalpole |
| 39 | Left-Pallidum | 81 | ctx-rh-temporalpole |
| 40 | Left-Hippocampus | 82 | ctx-rh-transversetemporal |
| 41 | Left-Amygdala | 83 | ctx-rh-insula |
| 42 | Left-Accumbens-area | 84 | Right-Cerebellum-Cortex |

## References

1. Bartolomei F, Lagarde S, Wendling F, McGonigal A, Jirsa V, Guye M, et al. Defining epileptogenic networks: contribution of SEEG and signal analysis. Epilepsia. 2017;58(7):1131–1147.

2. Proix T, Bartolomei F, Guye M, Jirsa VK. Individual brain structure and modelling predict seizure propagation. Brain. 2017;140(3):641–654.

3. Rosenow F, sLüders H. Presurgical evaluation of epilepsy. Brain. 2001;124(9):1683–1700.

4. Kuhlmann L, Lehnertz K, Richardson MP, Schelter B, Zaveri HP. Seizure prediction-ready for a new era. Nature Reviews Neurology. 2018;14(10):618–630.

5. Hashemi M, Vattikonda AN, Sip V, Diaz-Pier S, Peyser A, Wang H, et al. On the influence of prior information evaluated by fully Bayesian criteria in a personalized whole-brain model of epilepsy spread. PLoS computational biology. 2021;17(7):e1009129.

6. Makhalova J, Medina Villalon S, Wang H, Giusiano B, Woodman M, Bénar C, et al. Virtual epileptic patient brain modeling: Relationships with seizure onset and surgical outcome. Epilepsia. 2022;63(8):1942–1955.

7. Wang HE, Woodman M, Triebkorn P, Lemarechal JD, Jha J, Dollomaja B, et al. Delineating epileptogenic networks using brain imaging data and personalized modeling in drug-resistant epilepsy. Science Translational Medicine. 2023;15(680):eabp8982.

8. Jirsa V, Wang H, Triebkorn P, Hashemi M, Jha J, Gonzalez-Martinez J, et al. Personalised virtual brain models in epilepsy. The Lancet Neurology. 2023;22(5):443–454.

9. Jirsa VK, Proix T, Perdikis D, Woodman MM, Wang H, Gonzalez-Martinez J, et al. The Virtual Epileptic Patient: individualized whole-brain models of epilepsy spread. Neuroimage. 2017;145:377–388.

10. Hashemi M, Vattikonda A, Sip V, Guye M, Bartolomei F, Woodman MM, et al. The Bayesian Virtual Epileptic Patient: A probabilistic framework designed to infer the spatial map of epileptogenicity in a personalized large-scale brain model of epilepsy spread. NeuroImage. 2020;217:116839.

11. Kini LG, Bernabei JM, Mikhail F, Hadar P, Shah P, Khambhati AN, et al. Virtual resection predicts surgical outcome for drug-resistant epilepsy. Brain. 2019;142(12):3892–3905.

12. Sip V, Hashemi M, Vattikonda AN, Woodman MM, Wang H, Scholly J, et al. Data-driven method to infer the seizure propagation patterns in an epileptic brain from intracranial electroencephalography. PLoS computational biology. 2021;17(2):e1008689.

13. Vattikonda AN, Hashemi M, Sip V, Woodman MM, Bartolomei F, Jirsa VK. Identifying spatio-temporal seizure propagation patterns in epilepsy using Bayesian inference. Communications biology. 2021;4(1):1244.

14. Jha J, Hashemi M, Vattikonda AN, Wang H, Jirsa V. Fully Bayesian estimation of virtual brain parameters with self-tuning Hamiltonian Monte Carlo. Machine Learning: Science and Technology. 2022;3(3):035016.

15. Hashemi M, Vattikonda AN, Jha J, Sip V, Woodman MM, Bartolomei F, et al. Amortized Bayesian inference on generative dynamical network models of epilepsy using deep neural density estimators. Neural Networks. 2023;163:178–194.

16. Schirner M, Rothmeier S, Jirsa VK, McIntosh AR, Ritter P. An automated pipeline for constructing personalized virtual brains from multimodal neuroimaging data. NeuroImage. 2015;117:343–357. doi:10.1016/j.neuroimage.2015.03.055.

17. Fischl B. FreeSurfer. NeuroImage. 2012;62(2):774–781. doi:10.1016/j.neuroimage.2012.01.021.

18. Jenkinson M, Bannister P, Brady M, Smith S. Improved Optimization for the Robust and Accurate Linear Registration and Motion Correction of Brain Images. NeuroImage. 2002;17(2):825–841. doi:10.1006/nimg.2002.1132.

19. Tournier JD, Calamante F, Connelly A. Robust determination of the fibre orientation distribution in diffusion MRI: Non-negativity constrained super-resolved spherical deconvolution. NeuroImage. 2007;35(4):1459–1472. doi:10.1016/j.neuroimage.2007.02.016.

20. Tournier JD, Calamante F, Connelly A. Determination of the appropriatebvalue and number of gradient directions for high-angular-resolution diffusion-weighted imaging. NMR in Biomedicine. 2013;26(12):1775–1786. doi:10.1002/nbm.3017.

21. Tournier JD, Calamante F, Connelly A. Improved probabilistic streamlines tractography by 2nd order integration over fibre orientation distributions. In: Proceedings of the international society for magnetic resonance in medicine. vol. 18; 2010. p. 1670.

22. Desikan RS, Ségonne F, Fischl B, Quinn BT, Dickerson BC, Blacker D, et al. An automated labeling system for subdividing the human cerebral cortex on MRI scans into gyral based regions of interest. NeuroImage. 2006;31(3):968–980. doi:10.1016/j.neuroimage.2006.01.021.

23. Sanz-Leon P, Knock SA, Spiegler A, Jirsa VK. Mathematical framework for large-scale brain network modeling in The Virtual Brain. Neuroimage. 2015;111:385–430.

24. Jirsa VK, Haken H. Field Theory of Electromagnetic Brain Activity. Phys Rev Lett. 1996;77(5):960–963.

25. David O, Friston KJ. A neural mass model for MEG/EEG:: coupling and neuronal dynamics. NeuroImage. 2003;20(3):1743–1755. doi:10.1016/j.neuroimage.2003.07.015.

26. Deco G, Jirsa VK, Robinson PA, Breakspear M, Friston K. The dynamic brain: from spiking neurons to neural masses and cortical fields. PloS Comp Biol. 2008;4(8).

27. Cook BJ, Peterson AD, Woldman W, Terry JR. Neural Field Models: A mathematical overview and unifying framework. Mathematical Neuroscience and Applications. 2022;2.

28. Hashemi M, Hutt A, Sleigh J. Anesthetic action on extra-synaptic receptors: effects in neural population models of EEG activity. Frontiers in Systems Neuroscience. 2014;8:232.

29. Courtiol J, Guye M, Bartolomei F, Petkoski S, Jirsa VK. Dynamical Mechanisms of Interictal Resting-State Functional Connectivity in Epilepsy. Journal of Neuroscience. 2020;40(29):5572–5588. doi:10.1523/JNEUROSCI.0905-19.2020.

30. Lavanga M, Stumme J, Yalcinkaya BH, Fousek J, Jockwitz C, Sheheitli H, et al. The virtual aging brain: a model-driven explanation for cognitive decline in older subjects. bioRxiv. 2022;.

31. Yalcinkaya BH, Ziaeemehr A, Fousek J, Hashemi M, Lavanga M, Solodkin A, et al. Personalized virtual brains of Alzheimer’s Disease link dynamical biomarkers of fMRI with increased local excitability. medRxiv. 2023; p. 2023–01.

32. Jirsa VK, Stacey WC, Quilichini PP, Ivanov AI, Bernard C. On the nature of seizure dynamics. Brain. 2014;137(8):2210–2230.

33. El Houssaini K, Bernard C, Jirsa VK. The epileptor model: a systematic mathematical analysis linked to the dynamics of seizures, refractory status epilepticus, and depolarization block. Eneuro. 2020;7(2).

34. Saggio ML, Crisp D, Scott JM, Karoly P, Kuhlmann L, Nakatani M, et al. A taxonomy of seizure dynamotypes. Elife. 2020;9:e55632.

35. Haken H. Synergetics. Physics Bulletin. 1977;28(9):412.

36. Jirsa VK, Friedrich R, Haken H, Kelso JS. A theoretical model of phase transitions in the human brain. Biological cybernetics. 1994;71:27–35.

37. Hashemi M, Valizadeh A, Azizi Y. Effect of duration of synaptic activity on spike rate of a Hodgkin-Huxley neuron with delayed feedback. Physical Review E. 2012;85(2):021917.

38. Jafarian A, Zeidman P, Wykes RC, Walker M, Friston KJ. Adiabatic dynamic causal modelling. NeuroImage. 2021;238:118243.

39. Olmi S, Petkoski S, Guye M, Bartolomei F, Jirsa V. Controlling seizure propagation in large-scale brain networks. PLoS computational biology. 2019;15(2):e1006805.

40. Bancaud J, Angelergues R, Bernouilli C, Bonis A, Bordas-Ferrer M, Bresson M, et al. Functional stereotaxic exploration (SEEG) of epilepsy. Electroencephalography and clinical neurophysiology. 1970;28(1):85–86.

41. Durbin J, Koopman SJ. Time series analysis by state space methods. vol. 38. OUP Oxford; 2012.

42. Box GE, Jenkins GM, Reinsel GC, Ljung GM. Time series analysis: forecasting and control. John Wiley & Sons; 2015.

43. Hashemi M, Hutt A, Buhry L, Sleigh J. Optimal model parameter estimation from EEG power spectrum features observed during general anesthesia. Neuroinformatics. 2018;16:231–251.

44. Ionides EL, Bretó C, King AA. Inference for nonlinear dynamical systems. Proceedings of the National Academy of Sciences. 2006;103(49):18438–18443.

45. Izhikevich EM. Dynamical systems in neuroscience. MIT press; 2007.

46. Trentelman HL, Stoorvogel AA, Hautus M. Control theory for linear systems. Springer Science & Business Media; 2012.

47. Medaglia JD, Pasqualetti F, Hamilton RH, Thompson-Schill SL, Bassett DS. Brain and cognitive reserve: Translation via network control theory. Neuroscience & Biobehavioral Reviews. 2017;75:53–64.

48. Carver CS, Scheier MF. Control theory: A useful conceptual framework for personality–social, clinical, and health psychology. Psychological bulletin. 1982;92(1):111.

49. Scheid BH, Ashourvan A, Stiso J, Davis KA, Mikhail F, Pasqualetti F, et al. Time-evolving controllability of effective connectivity networks during seizure progression. Proceedings of the National Academy of Sciences. 2021;118(5):e2006436118.

50. Proix T, Jirsa VK, Bartolomei F, Guye M, Truccolo W. Predicting the spatiotemporal diversity of seizure propagation and termination in human focal epilepsy. Nature communications. 2018;9(1):1088.

51. Ghahramani Z, Hinton GE. Variational learning for switching state-space models. Neural computation. 2000;12(4):831–864.

52. Turner R, Deisenroth M, Rasmussen C. State-space inference and learning with Gaussian processes. In: Proceedings of the Thirteenth International Conference on Artificial Intelligence and Statistics. JMLR Workshop and Conference Proceedings; 2010. p. 868–875.

53. Frigola R, Chen Y, Rasmussen CE. Variational Gaussian process state-space models. Advances in neural information processing systems. 2014;27.

54. Archer E, Park IM, Buesing L, Cunningham J, Paninski L. Black box variational inference for state space models. arXiv preprint arXiv:151107367. 2015;.

55. Brunton SL, Proctor JL, Kutz JN. Discovering governing equations from data by sparse identification of nonlinear dynamical systems. Proceedings of the national academy of sciences. 2016;113(15):3932–3937.

56. Nassar J, Linderman SW, Bugallo M, Park IM. Tree-structured recurrent switching linear dynamical systems for multi-scale modeling. arXiv preprint arXiv:181112386. 2018;.

57. Koppe G, Toutounji H, Kirsch P, Lis S, Durstewitz D. Identifying nonlinear dynamical systems via generative recurrent neural networks with applications to fMRI. PLoS computational biology. 2019;15(8):e1007263.

58. Sip V, Hashemi M, Dickscheid T, Amunts K, Petkoski S, Jirsa V. Characterization of regional differences in resting-state fMRI with a data-driven network model of brain dynamics. Science Advances. 2023;9(11):eabq7547.

59. Penas DR, González P, Egea JA, Doallo R, Banga JR. Parameter estimation in large-scale systems biology models: a parallel and self-adaptive cooperative strategy. BMC bioinformatics. 2017;18:1–24.

60. Rodriguez-Fernandez M, Egea JA, Banga JR. Novel Metaheuristic for Parameter Estimation in Nonlinear Dynamic Biological Systems. BMC Bioinformatics. 2006;7(1):483.

61. Egea JA, Martí R, Banga JR. An evolutionary method for complex-process optimization. Computers & Operations Research. 2010;37(2):315–324.

62. de la Maza M, Yuret D. Dynamic hill climbing. AI Expert. 1994;9(3):26–31.

63. Penas DR, Henriques D, González P, Doallo R, Saez-Rodriguez J, Banga JR. A parallel metaheuristic for large mixed-integer dynamic optimization problems, with applications in computational biology. PLOS ONE. 2017;12(8):1–32.

64. Villaverde AF, Bongard S, Mauch K, Müller D, Balsa-Canto E, Schmid J, et al. A consensus approach for estimating the predictive accuracy of dynamic models in biology. Computer methods and programs in biomedicine. 2015;119(1):17–28.

65. Villaverde AF, Raimúndez E, Hasenauer J, Banga JR. Assessment of prediction uncertainty quantification methods in systems biology. IEEE/ACM transactions on computational biology and bioinformatics. 2022;.

66. Akiba T, Sano S, Yanase T, Ohta T, Koyama M. Optuna: A next-generation hyperparameter optimization framework. In: Proceedings of the 25th ACM SIGKDD international conference on knowledge discovery & data mining; 2019. p. 2623–2631.

67. Müller CL, Baumgartner B, Ofenbeck G, Schrader B, Sbalzarini IF. pCMALib: a parallel fortran 90 library for the evolution strategy with covariance matrix adaptation. In: Proceedings of the 11th Annual conference on Genetic and evolutionary computation; 2009. p. 1411–1418.

68. Penas DR, Banga JR, González P, Doallo R. Enhanced parallel Differential Evolution algorithm for problems in computational systems biology. Applied Soft Computing. 2015;33:86–99.

